# A general approach for selection of epitope-directed binders to proteins

**DOI:** 10.1101/2022.10.24.513434

**Authors:** Jie Zhou, Chau Q. Le, Yun Zhang, James A. Wells

**Affiliations:** Department of Pharmaceutical Chemistry, University of California San Francisco, San Francisco, CA, 94158, USA; Chan Zuckerberg Biohub, San Francisco, CA, 94158, USA; Department of Cellular and Molecular Pharmacology, University of California San Francisco, San Francisco, CA, 94158, USA

## Abstract

Directing antibodies to a particular epitope among many possible on a target protein is a significant challenge. Here we present a simple and general method for epitope-directed selection (EDS) using a differential phage selection strategy. This involves engineering the protein of interest (POI) with the epitope of interest (EOI) mutated using a systematic bioinformatics algorithm to guide the local design of an EOI decoy variant. Using several alternating rounds of negative selection with the EOI decoy variant followed by positive selection on the wild-type (WT) POI, we were able to identify highly specific and potent antibodies to five different EOI antigens that bind and functionally block known sites of proteolysis. Among these we developed highly specific antibodies that target the proteolytic site on the CUB domain containing protein 1 (CDCP1) to prevent its proteolysis allowing us to study the cellular maturation of this event that triggers malignancy. We generated antibodies that recognize the junction between the pro and catalytic domains for four different matrix metalloproteases (MMPs), such as MMP1, MMP3, and MMP9, that selectively block activation of each of these enzymes and impairs cell migration. We targeted a proteolytic epitope on the cell surface receptor, EPH Receptor A2, that is known to transform it from a tumor suppressor to an oncoprotein. We believe the EDS method greatly facilitates the generation antibodies to specific EOIs on a wide range of proteins and enzymes for broad therapeutic and diagnostic applications.

**Significance:** We have developed a highly efficient platform to facilitate the directed selection *in vitro* of antibodies to a wide range of functional epitopes on proteins. This method uses a bioinformatic program to guide mutations in the local site of interest to create a decoy antigen that can effectively remove antibodies not binding the site of interest by negative selection, followed by positive selection with the WT antigen to identify antibodies to the epitope of interest. We demonstrate the generality and versatility of this method by successfully producing functional antibodies to block specific proteolytically sensitive epitopes on five different proteins including enzymes important in cancer. The epitope-directed selection (EDS) approach greatly facilitates the identification of binders to specific sites of interest on proteins to probe function and as potential immunotherapeutics.

## Introduction

In the past two decades, therapeutic antibodies have gained prominence as the leading products in the biopharmaceutical market^*1*^. Most of these antibodies are chosen to block ligand-receptor interactions^*2*, *3*^, elicit agonistic activities^*4*^, modulate protein functions, or stabilize protein complexes^*1*^, by binding to a specific epitope on the protein of interest (POI). However, it remains challenging to directly identify binders to specific epitopes of interest (EOI) among many possible without extensive functional screening and characterization of antibodies produced by B-cell sorting from animal immunization^*5*^, or phage/yeast in vitro display technologies^*6*, *7*^.

There has been considerable effort to simplify identification of antibodies to a specific EOI on a POI. For example, animal immunization with linear peptides to EOIs has been used but is challenging because disordered peptides are poor mimics of the native 3D conformation, which is generally important for high affinity binding. The Wang group cleverly developed a method for generating epitope-directed antibodies using phage panning technology coupled with proximity photo-crosslinking, achieved by incorporating a noncanonical photoreactive amino acid into the EOI of the antigen^*8*^. However, this approach is limited to producing proteins with unnatural amino acids, creating a bump in the EOI, and incomplete photo-cross-linking which may produce non-specific binders. Another approach is to use epitope masking to block binding to undesirable sites using *in vitro* selection, but this requires characterization and isolation of multiple blocking antibodies^*9*^.

An area of emerging application for EOI antibodies is to block sites of proteolysis. Extracellular proteolysis has major functional consequences for activating and remodeling cell surface proteins and the extracellular matrix^*10-12*^. Alterations in proteolytic systems underlie multiple pathological conditions, including neurodegenerative disorders^*13*, *14*^, inflammatory diseases^*15*^, and cancers^*16*,*17*^. Preventing pericellular proteolysis specifically can yield valuable information regarding the role of proteolysis in regulating cellular activity, its implications in disease, and possible avenues for therapeutic intervention. For example, highly specialized antibodies antagonizing different Notch receptors by preventing their proteolytic activation has been very useful for dissecting the contributions of distinct Notch receptors to differentiation and disease^*18*^. Human tumors have the capability to inhibit natural killer (NK) and CD8^+^ T cell by avoiding NKG2D recognition through shedding of MICA/B from the cell surface. Antibodies or vaccines preventing MICA/B proteolytic cleavage on tumor cells lead to enhanced immune infiltration and anti-tumor responses^*19*^. Trastuzumab binding to HER2 blocks proteolytic cleavage of the extracellular domain of HER2, resulting in diminished levels of the more active p95–HER2 form of HER2.^*20*^

Here, we describe a general and simple technology that can be used for epitope-directed selection (EDS) using differential phage antibody selections *in vitro* (**Fig. 1a**). Differential phage selection has been powerfully applied to obtain highly specific antibodies for specific protein conformations^*21*^, isoforms^*22*^, and even small molecule complexes^*23*^. To extend differential selection for EDS we describe a design algorithm to produce a decoy antigen mutated in the EOI to deplete binders that recognize undesired epitopes from a Fab phage library before positive selection against the WT POI. After several rounds of iterative selection, only Fab phage that recognize the EOI with high affinity are enriched for further characterization. We validate the generality of EDS and apply it to effectively block the proteolysis of cell surface receptors in cancer (CDCP1, EphA2) and the proteolytic activation of pro-enzymes in cell migration (MMP1, MMP3, and MMP9). The EDS method allows deliberate selection of functional blockers to protein sites of interest.

**Figure 1.**
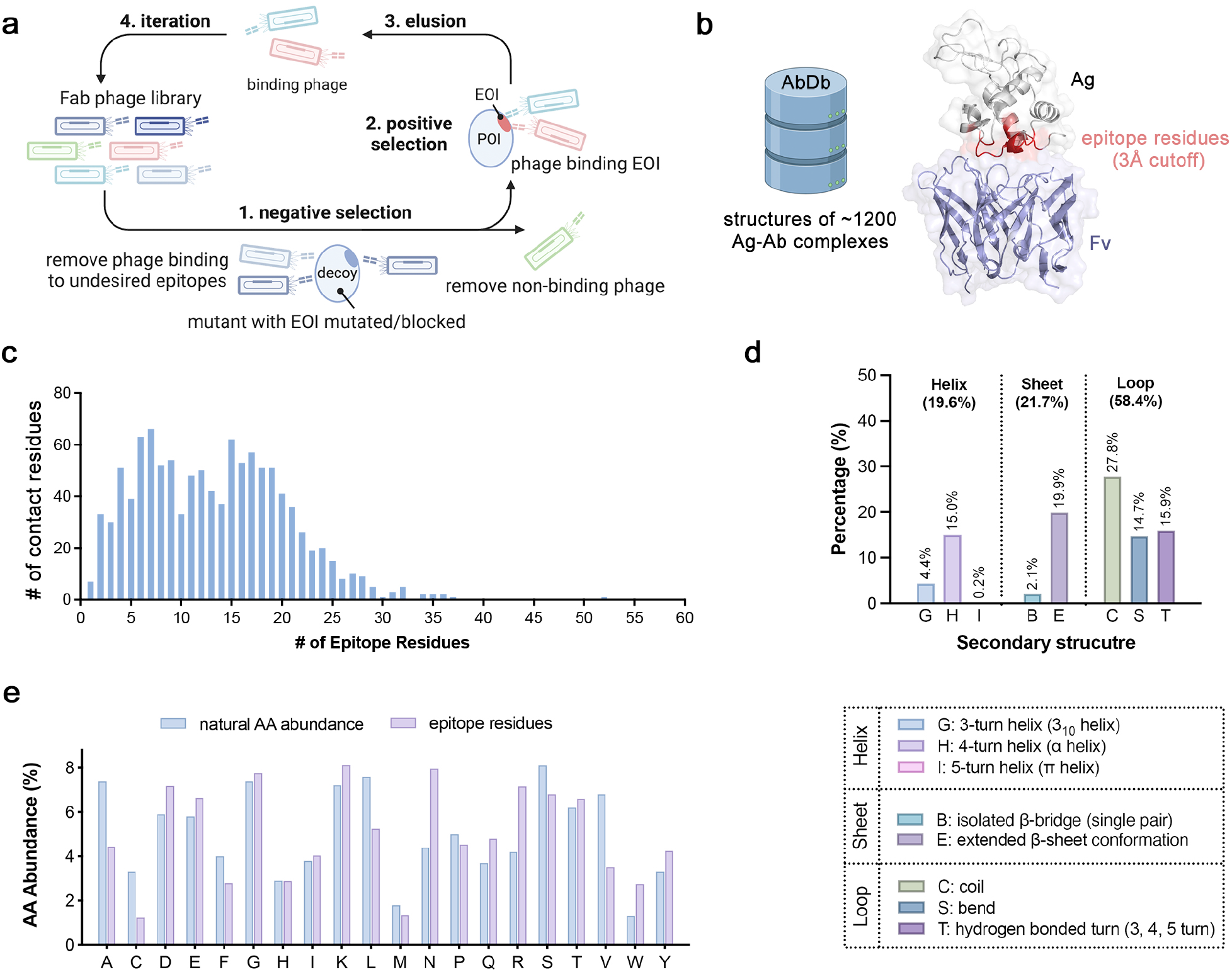
Strategy of epitope-directed selection using a Fab phage display. **(a)** EDS strategy to identify epitope-directed antibodies. A Fab phage library was pre-cleared with EOI mutated decoys prior to positive selection with WT POI. **(b)** Structural bioinformatics analysis of the epitope residues for 1200 antigen-antibody complexes using structures sourced from the AbDb database. (demo PDB: 1A2Y). **(c)** The frequency distribution for number of contact residues per epitope. (**d**) Antibodies contact antigen residues with higher frequency in loops (∼60%) than either helices (∼20%) or sheets (∼20%). The secondary structures containing the contact residues were calculated using the DSSP algorithm. **(e)** A comparison between the abundance of amino acids in the epitopes and their natural amino acid abundance.

## Results

### Structural bioinformatics guides the design of mutants/decoys for negative selection

We sought to develop a design protocol based on structural considerations, producing decoy antigens for negative selection with the smallest perturbations, in order to preserve the folded structure of the EOI decoy yet be disruptive to local binding at the EOI. There have been extensive bioinformatic studies to characterize CDR sequences and amino acid compositions of antibodies^*24-26*^, but less so on the antigen side^*27*^. We therefore conducted a structural bioinformatic analysis from the current ∼1200 antigen-antibody complex structures (sourced from AbDb^*28*^) to better understand typical number of contact residues, secondary structures within the epitope, and amino acid compositions (**Fig 1b-1e**). We find >90% of the antibodies contact 6-21 residues in a non-linear epitope on the antigen (**Fig 1c**). About 60% of the contacts are unstructured or in loop structures on the antigen and only 20% each for helix and sheet structures (**Fig 1d**). There was a bias for contacting large side chains (D, E, K, N, Q, R, W, Y) compared to small side chains (A, C, S), or some other hydrophobics (F, L, M) relative to their natural abundances (**Fig 1e**). In the final step, one needs to decide which amino acid to substitute. A common practice is to use alanine because it is intuitively most benign in that it simply deletes interactions, does not impose additional steric clash, and is tolerated in most secondary structures^*29*^. Based on our analysis we used three simple steps for designing EOI decoys for negative selection: (i) define the EOI on the antigen using PDB structures or AlphaFold models of the antigen, (ii) prioritize mutating large polar residues (D, E, K, N, Q, R), preferably in a loop, and (iii) mutate 4-6 of these to alanine or other small residues that generally are less disruptive to structure. As shown below, this approach leads to successful generation of decoys and antibodies to EOIs in five different POIs.

### Application of EDS to generate site-directed antibodies that block CDCP1 proteolysis

CDCP1 is a Type I single-pass membrane protein that is highly overexpressed in a variety of solid tumors and reported to afford a survival benefit to metastatic cells by preventing anoikis.^*30*, *31*^ It is proteolytically processed to generate an N-terminal and C-terminal fragment (NTF and CTF, respectively) that can activate signaling (**Fig. 2a**)^*32*^. We previously identified a nested set of three di-basic cleavage sites within six residues of each other between the NTF and CTF in pancreatic cell lines (**Fig. 2a**), and showed that the NTF stays tightly associated upon proteolysis.^*22*^

**Figure 2.**
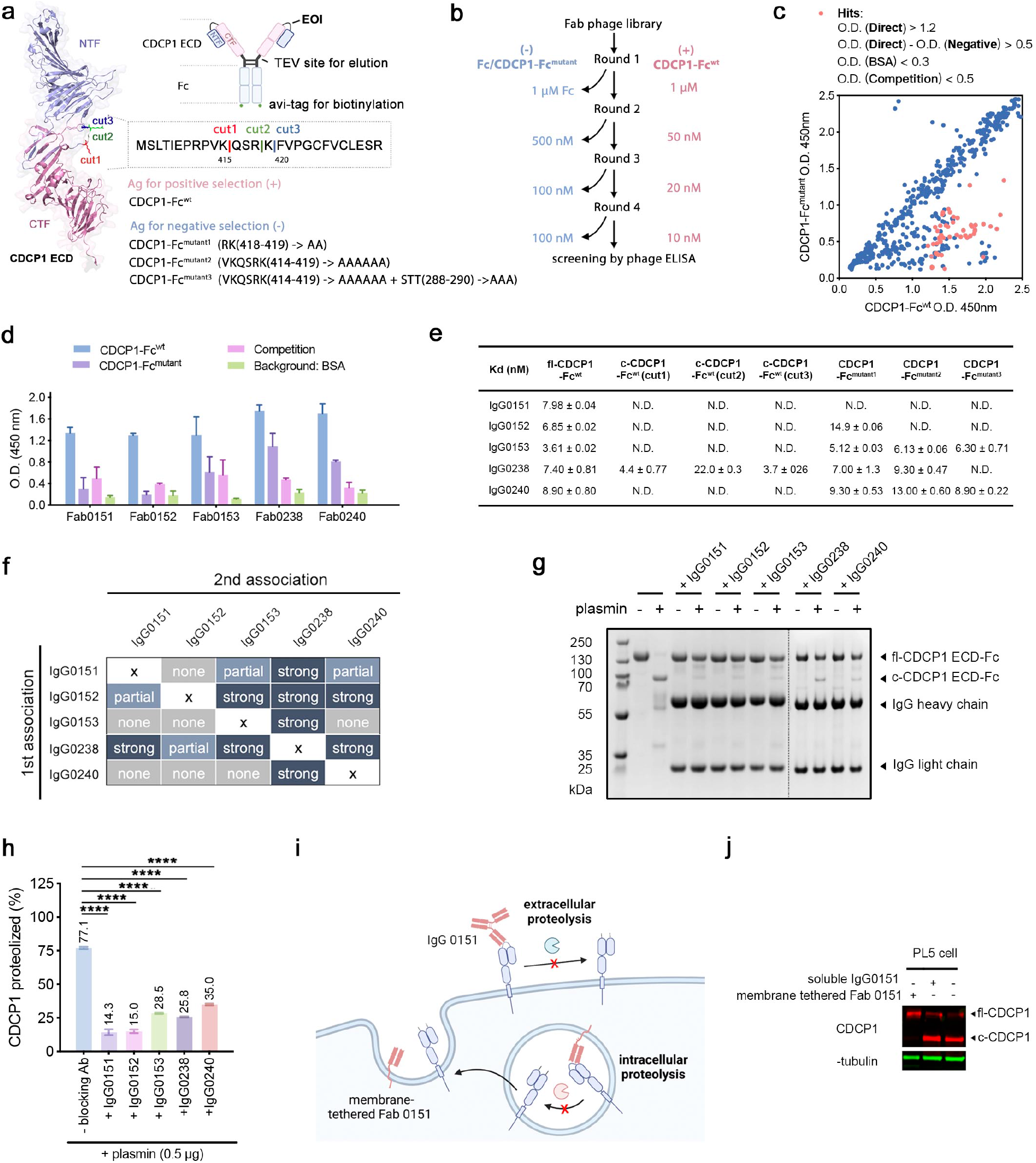
Generating EDS antibodies to block CDCP1 proteolysis. (**a**) Design of WT and EOI decoy antigens used for positive (+) and negative (-) selections, respectively, for EDS antibodies that block proteolysis of CDCP1. The structure of CDCP1 ECD domain was generated by AlphaFold2. (**b**) Workflow for the EDS strategy. (**c**) Screening of binding of Fab phage clones to CDCP1-Fc^wt^ and CDCP1-Fc^mutant1,2, or 3^ by single point phage ELISA. Hits (red dots) are defined as those that preferentially bind to CDCP1-Fc^wt^ over CDCP1-Fc^mutant1-3^, are easily competed off with soluble CDCP1-Fc^wt^, and exhibit minimum nonspecific binding. O.D. values for Direct, Negative, Competition, and BSA binding are defined in **Extended Data Fig. 1c**. (**d**) Five Fab hits, with unique sequences, were expressed, purified, and tested for binding to CDCP1-Fc^wt^ over their corresponding mutant used for negative selection, as well as being blocked by competition or tested for binding to BSA. (**e**) *In vitro* binding affinities for the five antibodies in IgG format to different CDCP1 constructs: uncleaved CDCP1-Fc^wt^, cleaved CDCP1-Fc^wt^ with three different cut forms, and CDCP1-Fc^mutant1, 2, or 3^. N.D. means below detection limit. (**f**) Heat map for epitope binning experiments by BLI revealed epitopes for the five antibodies. (**g**) Inhibition of plasmin cleavage of CDCP1 by the individual EDS antibodies shown by a reducing SDS-PAGE gel (1mM DTT) and (**h**) quantified by densitometry. **(i)** Strategy for testing whether proteolysis occurs outside the cell or during trafficking to the cell surface, comparing the level of cleavage of CDCP1 upon exogenous addition of the blocking antibody, IgG0151, or upon expression of a TM bound Fab0151, TM-Fab0151. **(j)** Result of the experiment in **(i)** showing exogenous IgG0151 has little impact on cleavage whereas TM-Fab0151 has marked effect.

We targeted the region containing the proteolytic sites (between R415-K420) to block proteolysis. First, we engineered three EOI decoys with increasing numbers of residues mutated. Next, we grafted the full-length extracellular domain (ECD) of WT CDCP1 onto a Fc domain containing a C-terminal Avi-tag for biotinylation and bead immobilization. A TEV cleavage site was also incorporated right before the hinge region between the ecto-domain of CDCP1 and Fc. This facilitated catch-and-release TEV elution of phage from the antigen bound to the bead (**Fig. 2a, Extended Data Fig. 1a**).^*33*^ Three EOI decoy mutants of CDCP1-Fc were generated by mutating the residues surrounding the proteolytic sites: CDCP1-Fc^mutant1^ (RK(418-419) -> AA); CDCP1-Fc^mutant2^ (VKQSRK(414-419) -> AAAAAA); and CDCP1-Fc^mutant3^ (VKQSRK(414-419) -> AAAAAA + STT(288-290) -> AAA). Each of these EOI decoys were expressed at levels comparable to the WT CDCP1-Fc suggesting that the mutations were well tolerated.

We conducted three rounds of differential selections by alternating negative selection with the EOI decoys followed by positive selection with the WT CDCP1-Fc (**Fig. 2b**). We utilized a synthetic diversity Fab phage library that has been extensively used for selecting Fabs to 100’s of targets.^*34*, *35*^ Prior to each round of selection, the phage pool was cleared with the mutant (CDCP1-Fc^mutant1, 2, or 3^) before positive selection with CDCP1-Fc^wt^, with the exception of the first-round clearance where Fc domain was used to remove Fc binders. After three to four rounds of iterative differential selection, there was significant enrichment for Fab phage that bound CDCP1-Fc^wt^ over the three EOI decoys (CDCP1-Fc^mutant1, 2, or 3^) characterized by standard phage tittering **(Extended Data Fig. 1b)**.

For each of the three selections, we characterized the binding for 96 phage clones to the CDCP1-Fc^wt^ and CDCP1-Fc^mutant1, 2, or 3^ by phage ELISA (**Extended Data Fig. 1c**). A group of Fab phage clones were identified to preferentially bind to the WT antigen over the EOI decoys with good affinity and low nonspecific binding (**Fig. 2c**). After Sanger sequencing, five unique clones were identified that preferentially bound to the CDCP1-Fc^wt^ over CDCP1-Fc^mutant1, 2, or 3^, and not to bovine serum albumin (BSA). Each of the five was effectively competed off with soluble CDCP1-Fc^wt^ (**Fig. 2d**).

We expressed all five clones as IgGs for further characterization against the uncleaved and three cleaved derivatives of CDCP1. A biolayer interferometry (BLI) experiment revealed that all the six IgGs bound specifically to uncleaved CDCP-Fc^wt^ with single-digit nM Kd values, but do not bind to most of the three proteolytically cleaved CDCP1 isoforms (**Fig. 2e, Extended Data Fig. 2**). Our previous studies indicated that cleaved and uncleaved CDCP1 have virtually identical conformations (**Fig. 2a**).^*22*^ This suggested that the EDS binders sit near and perhaps over the cleavage site, accounting for their lack of binding to the cleaved form. However, each of the six IgGs have subtle differences in binding specificity as indicated by differential binding affinities against the three different decoys, and epitope binning analysis (**Fig. 2f**). For example, IgG 0151, derived from the selection campaign using the decoy CDCP1-Fc^mutant1^ for negative selection, exhibits minimum binding to all three EOI decoys. In contrast, IgG 0152 from the selection using decoy CDCP1-Fc^mutant2^ for negative selection does not bind to CDCP1-Fc^mutant2^ as expected but does bind CDCP1-Fc^mutant1^ with only two-fold reduced affinity (Kd ∼15 nM). Moreover, IgG 0153 that was selected against CDCP1-Fc^mutant3^ strongly binds to both CDCP1-Fc^mutant1^ and CDCP1-Fc^mutant2^ decoys (KD value of 5 nM and 6 nM, respectively). These data indicates that R418 and K419 were required for IgG 0151 binding but not for IgG 0152 binding. IgG 0152 involves KQS (415-417) for binding while for IgG 0153, the VKQSRK site plays a negligible role in antibody binding. We further compared the binding of these five using standard epitope binning competition experiments by sequential binding on BLI (**Fig. 2f, Extended Data Fig. 3**). We find that some antibodies largely prevent binding of some of the others, suggesting they have the same or overlapping epitopes, while other antibodies do not block others, suggesting they have different epitopes. Overall, our studies indicate that the precise nature of the EOI decoys can finetune the selection of EDS antibodies, implying their significant utility in selecting for epitope-specific antibodies.

We next tested the ability of the EDS IgGs to prevent proteolysis by plasmin, which is known to cleave CDCP1 *in vitro*^*36*^. In the absence of antibody, about 77% of WT CDCP1-Fc (1 μg) was cleaved when treated with 0.5 μg plasmin for 1hr at room temperature. In contrast, the addition of any of the EDS IgGs at 1.5 molar equivalent resulted in only 13-35% cleaved CDCP1 in 1hr (**Fig. 2g, 2h)**, demonstrating the effectiveness of our antibodies in blocking proteolysis.

We were then interested in studying the biological relevance of these CDCP1 proteolysis-blocking antibodies. CDCP1 promotes cancer cell survival, growth, and metastasis because it is believed to inhibit anoikis. CDCP1 is known to participate in tumorigenic and metastatic signaling pathways, including the SRC/PKCδ, PI3K/AKT, Wnt, and RAS/ERK axes. Previous studies have shown that proteolysis of CDCP1 triggers the activation of CDCP1-CTF, leading to its phosphorylation by SRC kinase, homodimerization of CTF, and the formation of a phospho(p)-CDCP1-CTF/Src/PKCδ trimeric complex, which plays a crucial role in downstream signaling.^*32*^ Therefore, blocking CDCP1 function is a promising approach for cancer treatment. However, antibodies cannot enter the cell, so the effectiveness of a blocking EOI immunotherapy depends on the cellular location where proteolysis happens, either inside the cell during trafficking or outside the cell on the membrane (**Fig. 2i**). Although it is known that extracellular serine proteases with tryptic like specificity such as plasmin and uPA can cleave CDCP1, it is not known if this is relevant to the cellular landscape. To test this, we added IgG0151 exogenously to PL5 cells, a pancreatic cell line that predominantly displays cleaved CDCP1 on its surface. Remarkably, this did not prevent the proteolysis of CDCP1 as confirmed by Western Blot (WB) analysis. In striking contrast, when we overexpressed Fab0151 in PL5 cells as a membrane-tethered protein using a PDGF transmembrane domain (TM-Fab0151) it prevented virtually 100% of CDCP1 cleavage (**Fig. 2j**). These data strongly suggest that CDCP1 is processed by one or more proteases during trafficking in the ER and Golgi. These organelles house a number of di-basic pro-protein convertase whose specificity match the known cleavage sites. These data show the utility of EDS recombinant antibodies to block proteolysis and provide important tools for understanding proteolytic cell biology.

### Generating a novel side-directed inhibitory antibody to block MMP9 maturation

MMP9 plays a pathological role in a variety of inflammatory and oncology disorders and has long been considered an attractive therapeutic target for small molecules. However, it has been a challenging drug target due to its cross-reactivity with other MMP family members.^*37*^ MMP9, like the vast majority of proteases is produced as an inactive precursor, pro-MMP9. Inhibition is enforced by a conserved cysteine in the prodomain that ligands with the catalytic zinc atom in the active site. Proteolytic cleavage of the prodomain by other proteases releases the thiol-ligand constraint, leading to activation. The activation of pro-MMP9 can also be initiated by autolysis if the interaction between prodomain and catalytic core is disrupted by nonproteolytic means such as 4-aminophenylmercuric acetate (APMA), sodium dodecyl sulfate (SDS), or protein-protein interactions (**Fig. 3a**)^*38*^. Here we wanted to generate EDS antibodies that bind the junction between the prodomain and catalytic domain (**Fig. 3b**) to block pro-MMP9 activation as a novel class of MMP9 inhibitors.

To reduce the antigen size, we produced truncated forms of pro-MMP9 that deleted the fibronectin type II-like domain (Fn) and a C-terminal deletion of the hemopexin domain (Hpx) and fused them to an Fc (MMP9-pro-cat-Fc^wt^) (**Fig. 3b and 3c, Extended data Table 1**). We created two EOI decoys by mutating the residues immediately flanking the proteolytic activation site RF(106-107) -> AA and LGRFQT(104-109) -> AGAAAA, named MMP9-pro-cat-Fc^mutant1^ and MMP9-pro-cat-Fc^mutant2^, respectively (**Fig. 3b, 3c)**.

**Figure 3.**
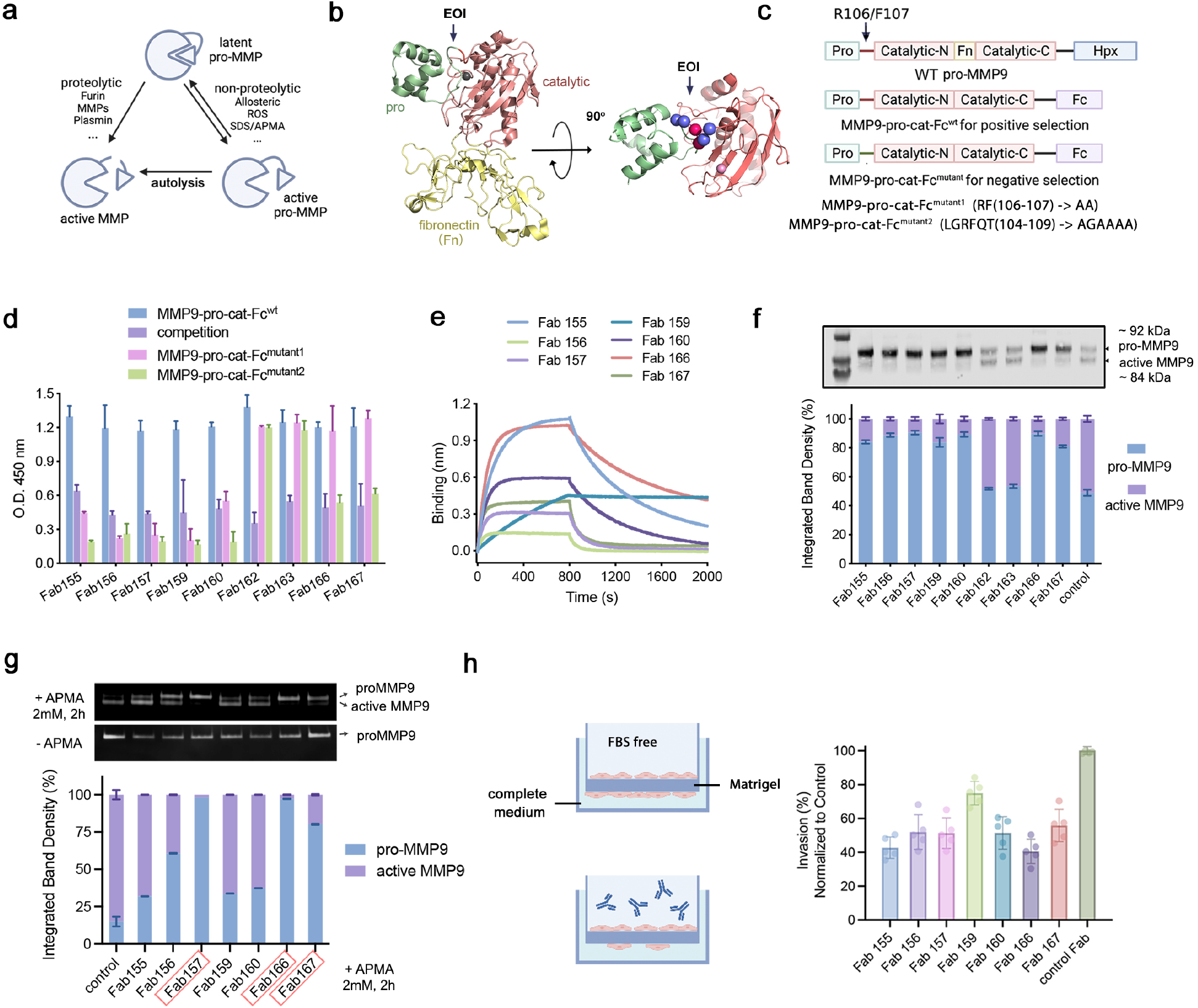
Generating EDS antibodies to block pro-MMP9 maturation as a novel class of inhibitors. (**a**) Diagram showing two mechanisms for pro-MMP activation/maturation. (**b**) Crystal structure of MMP9 pro-(catalytic-N)-Fn-(catalytic-C) domain. The catalytic domain of MMP9 consists of N and C-lobes with a fibronectin domain (Fn) in between. The EOI is highlighted in the top view. PDB: 1L6J. (**c**) The pro-MMP9 constructs designed for positive and negative phage panning. Truncated forms of pro-MMP9 were fused to a Fc with the Fn and Hemopexin (Hpx) domains truncated to reduce the antigen size. (**d**) Phage ELISA followed by Sanger sequencing identified seven out of nine unique clones that selectively bind MMP9-pro-cat-Fc^wt^ over at least one of the EOI mutants. (**e**) BLI confirmed the binding of the EDS Fabs (100 nM) to mammalian expressed MMP9-pro-cat-Fc^wt^ (25 nM) with varying affinities. (**f**) Seven out of nine EDS Fabs inhibit pro-MMP9 activation. HEK293T cells stably expressing pro-MMP9 (G100L) were treated with 25 ng/mL of each individual Fab overnight. Pro and active MMP9 in conditioned medium were visualized by WB analysis and then quantified using ImageJ. (**g**) Four of seven EDS Fabs block activation of endogenously secreted full-length pro-MMP9 induced by APMA ranging from 60 to over 95%. The pro-and active MMP9 forms were visualized by zymography analysis and quantified using ImageJ. (**h**) The cell invasion Matrigel assay (left diagram), where one can monitor migration from one side of a membrane to the other. Six of seven of the EDS Fabs inhibited cell migration of HT-1080 cells by 50% or greater (right bar graph).

The EDS campaign for pro-MMP9 was conducted as for CDCP1 and similarly generated significant phage titer enrichments based for the WT antigen over the EOI decoy. Phage ELISAs on 96 clones from each selection campaign was followed by Sanger sequencing, which identified seven out of nine unique Fab-phage clones that preferentially bound to the WT antigen over at least one of the two EOI mutants (**Fig. 3d, Extended Data Fig. 4a**). Notably, five of the seven Fabs did not bind either EOI mutant (155, 156, 157, 159, and 160), while two Fabs (166 and 167) retained binding to EOI Mutant1 which is less extensively modified than EOI Mutant2. BLI experiments of purified Fabs confirmed the selective binding to pro-MMP9 (**Fig. 3e, Extended Data Fig. 4b**). We also expressed and purified the truncated wild-type MMP9 with a C-terminal His-tag (MMP9-pro-cat-His^WT^) from *E. coli* inclusion bodies and confirmed the binding of these seven Fabs to the WT pro-MMP9 (**Extended Data Fig. 4c**).

A variant of secreted pro-MMP9 (G100L)^*39*^, one that weakens the interaction between the prodomain and the catalytic domain of pro-MMP9, is known to autoactivate when over-expressed in a stable HEK293T cell line. We tested the ability of exogenously added EDS Fabs to inhibit the auto-processing of this secreted pro-MMP9 variant by WB. Remarkably, seven out of nine EDS Fabs blocked pro-MMP9 processing by greater than 70%, with some over 90% (**Fig. 3f**) The two that did not block proteolysis (Fabs 162 and 163) retained binding to both EOI decoy mutants. Although we cannot rule out an allosteric mechanism, the simplest interpretation is that the inhibitory Fabs recognize proteolytic activation site on pro-MMP9, and the non-inhibitory ones do not.

4-Aminophenylmercuric acetate (APMA) promotes MMP9 activation through autolysis by chelating the inhibitory thiol, hence releasing the interaction between pro-domain and catalytic domain (**Fig. 3a**). We tested the ability of these Fabs to block activation on endogenously secreted full-length pro-MMP9 in the presence of APMA. This activation is almost completely blocked by addition of Fab157, 166, and 167, that were also shown above to be among the most potent inhibitors of autoactivation (**Fig. 3f and 3g**).

As MMP9 is thought to play a key role in tumor invasion and metastasis, we proceeded to test whether any of the seven EDS inhibitory antibodies would slow down metastasis of an invasive breast cancer line HT-1080 overexpressing MMP9. We conducted a Trans-well cell migration assay in the presence of EDS antibodies versus a nonbinding control antibody. The invasion or migration of HT-1080 cells through the Matrigel coated membrane was significantly slowed down (∼30-50%) depending on the EDS inhibitory antibody (**Fig. 3h**). The most potent Fabs (157, 166 and 167) were also those that scored highest in inhibiting the processing and activation assays, **Fig. 3f** and **Fig.3g**, respectively. Some of the Fabs showed interesting differences in the assays, such as 155 and 160 which were potent in inhibiting autoproteolysis and cell migration but far less so in the APMA activation assay. Future structural studies will be important to characterize the binding sites for all these Fabs to reveal direct versus allosteric means of inhibition. Overall, the MMP9 EDS antibody inhibitors offer a unique mechanism to target catalytic sites compared to traditional MMP inhibitors and could provide additional sites to perturb this important pro-enzyme.

### Generalizing differential phage selection approach to other MMPs

To further demonstrate the generality of the EDS approach, we developed highly selective blockers of the proteolytic activation to two other MMPs (i.e., MMP1, MMP3) that play roles in multiple cancers and are thought to be promising cancer targets. Like MMP9, these MMPs are all activated by proteolysis of a loop structure (**Fig. 4a, 4b**). Due to their common active sites, there has been a lack of selective small molecule inhibitors for individual MMPs, hampering further biological and drug discovery studies.

**Figure 4.**
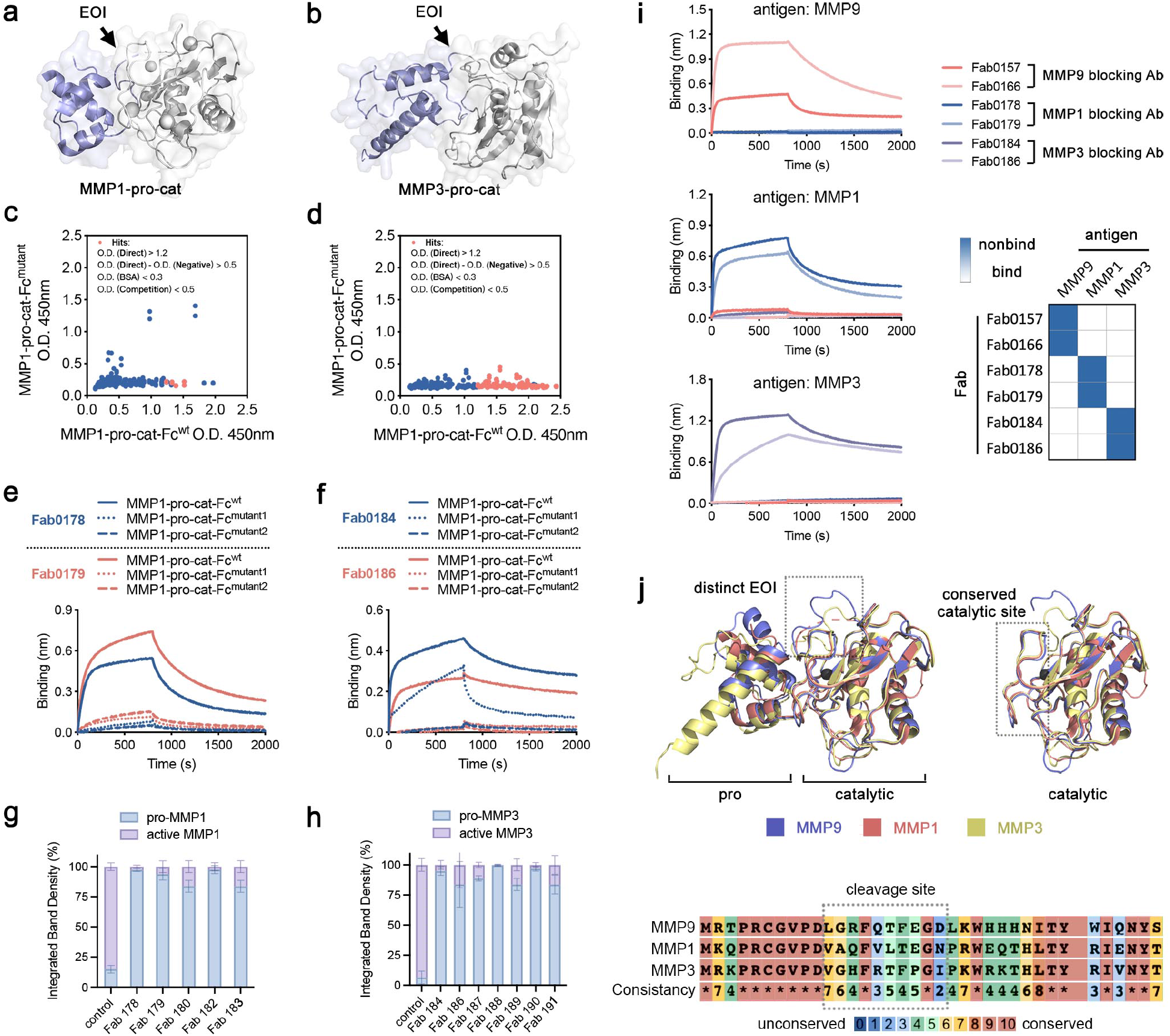
Generalizing EDS approach to other MMPs. (**a, b**) Crystal structure of pro-catalytic domains MMP1 and MMP3. EOIs are indicated by an arrow. PDB: 1SU3 for MMP1, 1SLM for MMP3. (**c, d**) 96 clones were characterized by phage ELISA for each campaign. A remarkably high proportion of clones (red) were found to preferentially bind WT antigens over their EOI variants. (**e, f**) BLI characterization of selective clones from each selection campaign in Fab format binding to WT pro-MMP antigens. These Fabs selectively bind to WT antigens over at least one of the two EOI decoys. (**g, h**) pro-MMP activation of MMP1 and MMP3 was inhibited by different EDS Fabs. Pro-MMPs and active MMPs were visualized by SDS-PAGE and quantification was conducted ImageJ. (**i**) The EDS Fabs show high specificity as they do not bind other MMP family members. (**j**) Structure and sequence alignments of pro or active MMP1, 3, and 9. PDB:1SU3 for MMP1, 1SLM for MMP3, 1L6J for MMP9. Sequence alignment was generated by PRALINE^*45*^.

We produced EOI decoys for each pro-MMP as we did to pro-MMP9, generating the MMP-pro-cat Fc fusion (MMP1, 3-pro-cat-Fcwt) (**Extended Data Table 1**). We conducted the EDS campaigns as we did for CDCP1 and pro-MMP9 and observed a robust enrichment for the WT antigens over the decoys for all three pro-MMP targets. Phage ELISA on 96 clones from each selection campaign followed by Sanger sequencing identified at least five unique Fab clones for each that preferentially bind to the WT antigens over at least one of the EOI decoy mutants (**Fig. 4c, 4d**). Fabs were expressed and purified, and BLI experiments confirmed the selective binding to the wild-type antigens (**Fig. 4e, 4f**) with impaired or minimum binding to the EOI decoys (**Extended Data Fig. 5, 6**). Finally, we validated that the EDS antibodies could efficiently inhibit proteolytic activation of their respective pro-MMP1 and 3 *in vitro* (**Fig. 4g, 4h**). The EDS antibodies selectively target the parent pro-MMP they were selected to bind and not the other family members demonstrating high specificity (**Fig. 4i**). This selectivity resulted from the unique sequences and structural features of the EOIs present on MMP1, MMP3, and MMP9 (**Fig. 4j, Extended Data Fig. 7**). It is noteworthy that the catalytic domains of the active isoforms of these MMPs are highly conserved, which typically prevents selective targeting. Thus, our findings underscore the significance of strategy with the EDS antibodies to identify selective binders that can block proteolytic activation.

### Generalizing differential phage selection approach to the cell surface receptor EphA2

Next, we wanted to expand the EDS selection strategy to the cell surface receptor EphA2. EphA2 can be cleaved by the cell surface protease, MT1-MMP, between the Fibronectin type-III domain 1 and 2 (Fn-III 1 and Fn-III 2, respectively). This converts EphA2 from a tumor suppressor to an oncogene (**Fig. 5a, Extended Data Fig. 8)**^*40-42*^. We produced an EOI decoy by mutating the SINQT sequence (433-437) to AAAAA in the loop connecting Fn-III domain-1 and Fn-III domain-2. We refer to this variant as EphA2-Fn-III-Fcmutant (**Fig 5a**). EDS was carried out as for the other four proteins above. Again, we found significant enrichment for the WT versus EOI decoy, and significant binding by Fab Phage ELISA (**Fig. 5b, 5c**). BLI experiments identified Fabs preferentially binding to WT over mutant (**Fig. 5d, Extended Data Fig. 9**).

**Figure 5.**
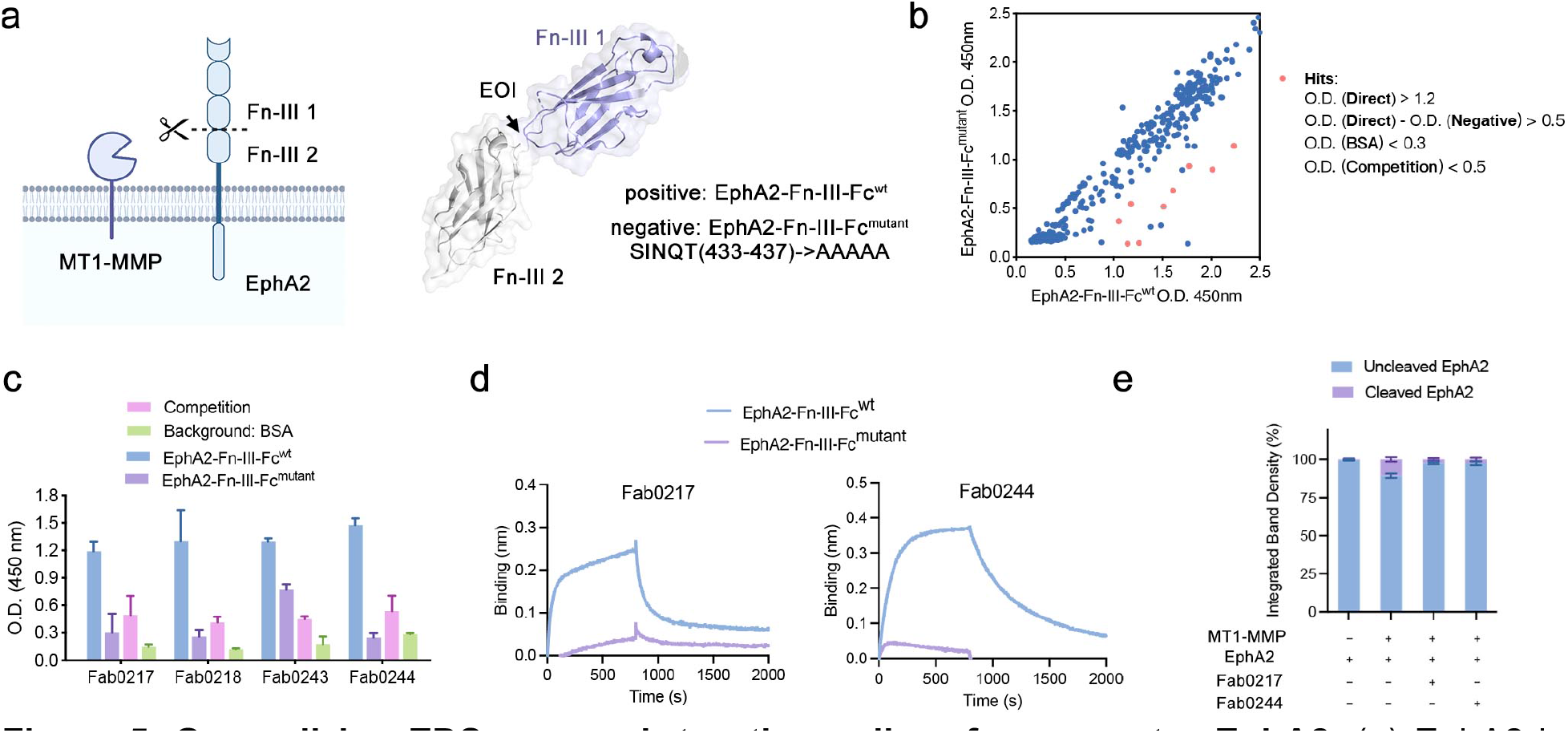
Generalizing EDS approach to other cell surface receptor EphA2. (**a**) EphA2 is known to be cleaved by the cell surface protease MT1-MMP between the Fibronectin type-III domain 1 and 2. (**b, c**) 288 clones were characterized by phage ELISA. A remarkably high proportion of clones (red) were found to preferentially bind WT antigens over the mutant. These Fab phage were Sanger sequenced to identify unique clones (**c**). (**d**) BLI characterization of selective clones from the selection campaign in Fab format binding to WT antigen. (**e**) Proteolysis of EphA2 by MT1-MMP is inhibited by EDS Fabs.

To test the ability of the EDS antibodies to inhibit cleavage of EphA2 by MT1-MMP, we stably expressed both EphA2 and MT1-MMP (MT1-WT) in HEK293T cells. Cell lysates were analyzed by Western blotting using a commercially available mAb and detected the known 130- and 100-kDa forms, which correspond to glycosylated (mature) and unglycosylated (immature) forms of EphA2 (**Fig. 5e**). An anti-MT1-MMP mAb detected both the overexpressed MT1-MMP (55 and 60 kDa) in the cells. As expected, overexpression of MT1-MMP increased the amount of two smaller EphA2-related cleaved fragments (60 and 65 kDa), as compared with cells expressing EphA2-myc alone. Treatment of cells with the EphA2 blocking EDS antibodies Fab0238 and Fab0244 abrogated cleavage of the two fragments (**Fig. 5e**). Thus, in contrast to the results with CDCP1, the proteolysis of EphA2 by MT1-MMP could be blocked by exogenous addition of an inhibitory EDS antibody.

## Discussion

Proteolysis is probably the most important regulated post-translational modification in the extracellular space. Our data suggest that EDS is a generalizable platform for identifying antibodies to specific EOIs on proteins. A key feature of this method is to create a decoy antigen by mutating the EOI to deplete the Fab phage library of any binders that are off target. We provide a simple design strategy based on structural bioinformatics to mutate anywhere from two to six large, predominantly polar side chains in loop regions to alanine (**Fig. 1**). The generality and versatility of this approach was demonstrated by the successful generation of functional antibodies that block proteolysis to defined epitopes on five different protein targets with distinct structures and biological roles. These recombinant antibodies are utilized to selectively inhibit protease activation by preventing pro-peptide cutting. Moreover, these antibodies help elucidate the subcellular location and secretory sequence of such proteolysis events. By testing the impact of exogenously added blocker versus intracellular expressed blocker, we provided strong evidence that CDCP1 is processed in transit to the cell membrane, whereas EphA2 appears to be processed on the cell surface. We believe these antibodies have the potential to be transformed into novel disease therapeutics and are valuable tools to study protein functions in various signaling pathways.

The differential phage selection strategy has great adaptability and flexibility. The selections generated potent and selective binders, from mutating only two residues in the EOI to a broadened mutation range of 6-8 residues. It was possible to tune the stringency of the selection by lowering the EOI decoy antigen concentration in the negative selection step. In this way, we found antibodies that bind narrowly or broadly to the EOI. We believe the EDS platform could be used to target specific loops on multi-pass membrane spanning proteins, such as an ion channels or G protein–coupled receptors. In such case, on-cell phage panning could be coupled with this differential phage selection to get site-directed antibodies that would inhibit or stimulate the signaling transduction. It should also be possible to generate catalytic site-specific antibodies for an enzyme by slightly modifying this protocol. For example, a negative selection can be performed with the enzyme pre-bound to any non-specific inhibitors targeting catalytic sites, then the cleared library is carried to positive selection with apo enzyme, allowing for enrichment of binders specifically targeting the catalytic site. Finally, the same principle would be used in other combinatorial selection technologies, including yeast display-based antibody generating pipeline.

To summarize, we have developed a general approach to select and identify epitope-directed antibodies to virtually any surface epitope. We demonstrate the generality and versatility of this method by producing binders blocking proteolytic activation of a panel of MMPs and two cell surface receptors, CDCP1 and EphA2. While the theme of this manuscript is inhibition of proteolytic activation, this strategy could be expanded to generate functional epitope-directed binders that block protein post-translational modifications (PTMs) beyond proteolysis, block protein-protein interaction sites (PPIs), elicit agonistic or antagonistic activities on receptors, stabilize protein complexes, or trap specific protein conformations. We believe the EDS approach enables the generation of binders to the sites of interest on a wide variety of proteins for broad therapeutic and diagnostic applications. In principle, the EDS approach need not be limited to proteins. It could be used for any target that can bind an antibody, for which a decoy target can be constructed. Targets could include peptides, small molecules, natural products, carbohydrates, nucleic acids or other synthetic or biomaterials, allowing for the discovery of new site-specific binders on these important biomolecules.

## Materials and Methods

### Structure bioinformatics

Secondary structures for residues in the epitope involved in binding were analyzed using an in-house informatics pipeline written in R. Scripts and are available for download at github. The antigen-Ab complex structures used for this analysis were downloaded from AbBD^*28*^. A DSSP algorithm was used to obtain the exact secondary structure information have been described^*43*^.

### Cloning, protein expression, and purification

Plasmids encoding WT or decoy CDCP1, MMP1, MMP3, MMP9 and EphA2 as Fc fusions or the heavy and light chains of IgGs, were generated by Gibson cloning into pFUSE vector (InvivoGen). Fab genes were PCR amplified from the Fab phagemid and subcloned into pBL347 Fab expression vector. Plasmids for stable cell line construction were generated by Gibson cloning into pCDH-EF1-CymR-T2A-Neo (System Bioscience) vector. The plasmid for *E coli* expression of MMP9-pro-cat-His^wt^ was generated by Gibson cloning into pET vector. Sequences of all plasmids were confirmed by Sanger sequencing.

Antigens for differential phage panning or antibody binding in IgG format were expressed by transient transfection of BirA-Expi293 cells (Life Technologies) with plasmids encoding CDCP1, MMPs and EphA2 as Fc fusions, or light/heavy chains of IgG. The ExpiFectamine 293 transfection kit (Life Technologies) was used for transfections as per manufacturer’s instructions. Cells were incubated for 4-5 days at 37°C in 5% CO2 at 125 rpm before the supernatants were harvested. Proteins were purified by Protein A affinity chromatography (Fc-fusions and IgGs) and assessed for quality and integrity by SDS-PAGE.

MMP9-pro-cat-His^wt^ (E coli expression) was expressed in BL21(DE3) (Thermo Fisher). Cells were grown in 2xYT at 37°C and expression was induced with 0.5 mM isopropyl β-D-1-thiogalactopyranoside (IPTG) for 3 hours. The inclusion bodies were harvested, and the protein was refolded as previously described^*39*^.

To express Fabs, C43 (DE3) Pro^+^ E. coli transformed with the Fab expression plasmid were grown in TB autoinduction media at 37°C for 6 hrs, then switched to 30°C for 16–18 hrs^*44*^. Cells were harvested by centrifugation (6000xg for 20 min) and lysed with B-PER Bacterial Protein Extraction Reagent (Thermo Fischer) supplemented with DNAse I (GoldBio). Lysate was incubated at 60°C for 20 min and cleared by centrifugation at 14,000xg for 30 min. Clarified supernatant was passed through a 0.45 μm syringe filter. Fabs were purified by Protein A affinity chromatography on an AKTA Pure system. Fab purity and integrity were assessed by SDS-PAGE.

### Phage selection

The phage selection protocol was adapted from a previously published study^*44*^. Briefly, selections were performed using biotinylated Fc fusion antigen captured on SA-coated magnetic beads (Promega). Prior to each selection, the phage pool was incubated with increasing concentrations of biotinylated negative selection antigen captured on streptavidin beads to deplete the library of any binders to the undesired epitope. Four rounds of selection were performed with decreasing amounts of positive selection antigens (**Fig. 2a**) captured on the SA beads. We employed a “catch and release” strategy, where bound Fab phage were eluted from the magnetic beads by the addition of 2 μg/mL of TEV protease. Individual phage clones from the third or fourth round of selection were analyzed for binding by phage ELISA.

### Phage ELISA

Phage ELISAs were performed according to standard protocols (**Extended Data Fig. 1c**). Briefly, 384-well Maxisorp plates were coated with NeutrAvidin (10 μg/mL) overnight at 4°C and subsequently blocked with PBS + 0.2% BSA for 1 hr at r.t. 20 nM of biotinylated positive/negative selection antigens were individually captured on the NeutrAvidin-coated wells for 30 min followed by the addition of phage supernatants diluted 1:5 in PBSTB for 30 min. The competition wells with captured biotinylated positive selection antigen were incubated with 1:5 diluted phage in the presence of 20 nM soluble positive selection antigen. Bound phage were detected using a horseradish peroxidase (HRP)-conjugated anti-M13 phage antibody (GE Lifesciences).

### CDCP1 proteolysis blocking assay

Uncleaved CDCP1 ECD Fc fusion (CDCP1-Fc^wt^) (1 μg) was treated with 0.5 μg of plasmin in 10 μL PBS buffer for 1 hour at room temperature, with or without 1.5 molar equivalent IgGs. The samples were loaded on an SDS-PAGE gel to assess the proteolysis of CDCP1.

### Mammalian Cell Culture

All the cell lines used for this study were originally purchased from the American Type Culture Collection (ATCC). The HEK293T, PL5, HT-1080, or HEK293T Lenti-X cell lines was cultured in DMEM + 10% FBS + 1% Pen/Strep. MCF-7 cells were cultured in McCoy’s 5A + 10% FBS + 1X Pen/Strep. Cell line identities were authenticated by morphological inspection. All cell lines were previously authenticated and tested for mycoplasma.

### Lentiviral cell line construction

The HEK293T stable cell line overexpressing MMP9 (G100L) and HT-1080 stable cell overexpressing EphA2 and MT1-MMP1 were generated by lentiviral transduction. To produce virus, HEK293T Lenti-X cells were transfected with a mixture of second-generation lentiviral packaging plasmids at ∼70% confluence. FuGene HD (Promega) was used for transfection of the plasmids using 3 μg DNA (1.35 μg pCMV delta8.91, 0.15 μg pMD2-G, 1.5 μg pCDH vectors encoding gene of interest) and 7.5 μL of FuGene HD per well of a six-well plate. Media was changed to complete DMEM after 6 hrs of incubation with transfection mixture. The supernatant was harvested and cleared by passing through a 0.45 μm filter 72 hrs post transfection. Cleared supernatant was added to target HEK293T or HT-1080 WT cells (∼1 million cells per mL) with 8 μg/mL polybrene and cells were centrifuged at 1000xg at 33°C for 2 hrs. Cells were then incubated with viral supernatant mixture overnight before the media was changed to fresh complete DMEM. Cells were expanded for a minimum of 48 hrs before they were grown in drug selection media. Drug selection for stable cell lines was started by the addition of 2 μg/mL puromycin or 500 μg/mL zeocin. Following at least 72 hrs of incubation in drug containing media, cells were assay or analyzed.

### Bio-layer interferometry (BLI) experiments

BLI experiments were performed using an Octet RED384 instrument (ForteBio). Biotinylated proteins were immobilized on a streptavidin (SA) biosensor and His-tagged proteins were immobilized on a Ni-NTA biosensor. Different concentrations of analyte in PBS pH 7.4 + 0.05% Tween-20 + 0.2% BSA (PBSTB) were used as analyte. K_D_ values were calculated from a global fit (1:1) of the data using the Octet RED384 software.

### Western Blot

Cell lysate or conditioned medium were harvested and supplemented with protease inhibitor cocktail (Roche). After centrifugation at 14,000xg for 30 min at 4°C, the supernatant was then transferred into a new tube and protein concentration was determined by BCA assay (ThermoFisher). Immunoblotting was performed using MMP-9 (D6O3H) XP® Rabbit mAb (primary, Cell Signaling, 13667T), and IRDye 680RD Goat anti-Mouse (secondary, LiCOR, 925– 68070).

### Zymography analysis

Conditioned medium was harvested and then concentrated 10-fold using a 10K cutoff spin concentrator (EMD Millipore). 4X non-reducing sample buffer (BioRad) was added to the sample, which was then loaded to the Novex™ 10% Zymogram Plus (Gelatin) Protein Gel. The gel was run at 200V until good band separation was achieved. The gel was incubated in Invitrogen Novex Zymograph renaturing buffer (Thermo Scientific) for 30 min, followed by Invitrogen Novex Zymograph developing buffer (Thermo Scientific) overnight. The gel was then stained in InstantBlue™ Coomassie Stain (Abcam) and imaged with a BioRad gel imager.

### Cell migration assay

The Trans-well cell invasion assay was conducted using Costar 6.5 mm Trans-well kit (8.0 μm Pore, Corning) per manufactory instructions. In brief, the insert was pre-coated with Matrigel (Thermo Scientific) in a 37°C incubator for 15-30 minutes to form a thin gel layer. A cell suspension (100 μL, 10^6^ cells/mL, FBS free) was added on top of the Matrigel coating to simulate invasion through the extracellular matrix with or without individual inhibitory Fabs (50 μg/mL). Complete medium was added to the bottom wells. Cells that migrated through the membrane were quantified using Crystal Violet (Fisher Scientific).

### Reporting Summary

Further information on research design is available in the Nature Research Reporting Summary linked to this article.

## Data availability

All data are available in the main text or the supplementary material. Any other data relating to this study are available from the corresponding authors on reasonable request.

## Acknowledgments

We thank members of the Wells Lab for helpful discussion and input, in addition to members of the Recombinant Antibody Network for feedback on this project. JZ is supported by a National Institutes of Health (NIH) National Cancer Institute F32 5F32CA236151. JAW acknowledges funding from NIH NCI 1P41CA196276, CA191018, and NIH GM097316 and commercial funding from Bristol Myers Squibb.

## Reference

1. Weiner, L. M., Surana, R., and Wang, S. Z. Monoclonal antibodies: versatile platforms for cancer immunotherapy, Nature Reviews Immunology 10, 317–327 (2010).

2. Mahoney, K. M., Rennert, P. D., and Freeman, G. J. Combination cancer immunotherapy and new immunomodulatory targets, Nature Reviews Drug Discovery 14, 561–584 (2015).

3. Topalian, S. L., Taube, J. M., Anders, R. A., and Pardoll, D. M. Mechanism-driven biomarkers to guide immune checkpoint blockade in cancer therapy, Nature Reviews Cancer 16, 275–287 (2016).

4. Waldmann, T. A., and O’Shea, J. The use of antibodies against the IL-2 receptor in transplantation, Current Opinion in Immunology 10, 507–512 (1998).

5. Green, L. Transgenic mouse strains as platforms for the successful discovery and development of human therapeutic monoclonal antibodies, Current drug discovery technologies 11 1, 74–84 (2014).

6. Winter, G., Griffiths, A. D., Hawkins, R. E., and Hoogenboom, H. R. Making antibodies by phage display technology Annual Review of Immunology 12, 433–455 (1994).

7. Gai, S. A., and Wittrup, K. D. Yeast surface display for protein engineering and characterization, Current Opinion in Structural Biology 17, 467–473 (2007).

8. Chen, L. X., Zhu, C. Y., Guo, H., Li, R. T., Zhang, L. M., Xing, Z. Z., Song, Y., Zhang, Z. H., Wang, F. P., Liu, X. F., et al. Epitope-directed antibody selection by site-specific photocrosslinking, Science Advances 6 (2020).

9. Ditzel, H. J. Rescue of a broader range of antibody specificities using an epitope-masking strategy, Antibody Phage Display: Methods and Protocols, 179–186 (2002).

10. Werb, Z. ECM and cell surface proteolysis: Regulating cellular ecology, Cell 91, 439–442 (1997).

11. Kessenbrock, K., Plaks, V., and Werb, Z. Matrix Metalloproteinases: Regulators of the Tumor Microenvironment, Cell 141, 52–67 (2010).

12. Sato, H., Takino, T., Okada, Y., Cao, J., Shinagawa, A., Yamamoto, E., and Seiki, M. A Matrix Metalloproteinase Expressed On The Surface Of Invasive Tumor-Cells, Nature 370, 61–65 (1994).

13. Yan, R. Q., Bienkowski, M. J., Shuck, M. E., Miao, H. Y., Tory, M. C., Pauley, A. M., Brashler, J. R., Stratman, N. C., Mathews, W. R., Buhl, A. E., et al. Membrane-anchored aspartyl protease with Alzheimer’s disease beta-secretase activity, Nature 402, 533–537 (1999).

14. Vassar, R., Bennett, B. D., Babu-Khan, S., Kahn, S., Mendiaz, E. A., Denis, P., Teplow, D. B., Ross, S., Amarante, P., Loeloff, R., et al. beta-secretase cleavage of Alzheimer’s amyloid precursor protein by the transmembrane aspartic protease BACE, Science 286, 735–741 (1999).

15. Tedgui, A., and Mallat, Z. Cytokines in atherosclerosis: Pathogenic and regulatory pathways, Physiological Reviews 86, 515–581 (2006).

16. Friedl, P., and Alexander, S. Cancer Invasion and the Microenvironment: Plasticity and Reciprocity, Cell 147, 992–1009 (2011).

17. Friedl, P., and Wolf, K. Tumour-cell invasion and migration: Diversity and escape mechanisms, Nature Reviews Cancer 3, 362–374 (2003).

18. Wu, Y., Cain-Hom, C., Choy, L., Hagenbeek, T. J., de Leon, G. P., Chen, Y., Finkle, D., Venook, R., Wu, X., Ridgway, J., et al. Therapeutic antibody targeting of individual Notch receptors, Nature 464, 1052–1057 (2010).

19. Badrinath, S., Dellacherie, M. O., Li, A., Zheng, S., Zhang, X., Sobral, M., Pyrdol, J. W., Smith, K. L., Lu, Y., Haag, S., et al. A vaccine targeting resistant tumours by dual T cell plus NK cell attack, Nature 606, 992–998 (2022).

20. Molina, M. A., Codony-Servat, J., Albanell, J., Rojo, F., Arribas, J., and Baselga, J. Trastuzumab (herceptin), a humanized anti-Her2 receptor monoclonal antibody, inhibits basal and activated Her2 ectodomain cleavage in breast cancer cells, Cancer research 61, 4744–4749 (2001).

21. Gao, J., Sidhu, S. S., and Wells, J. A. Two-state selection of conformation-specific antibodies, Proceedings of the National Academy of Sciences of the United States of America 106, 3071–3076 (2009).

22. Lim, S. A., Zhou, J., Martinko, A. J., Wang, Y.-H., Filippova, E. V., Steri, V., Wang, D., Remesh, S. G., Liu, J., Hann, B., et al. Targeting a proteolytic neo-epitope of CUB-domain containing protein 1 in RAS-driven cancer, bioRxiv, 2021.2006.2014.448427 (2021).

23. Hill, Z. B., Martinko, A. J., Nguyen, D. P., and Wells, J. A. Human antibody-based chemically induced dimerizers for cell therapeutic applications, Nature Chemical Biology 14, 112-+ (2018).

24. Koide, S., and Sidhu, S. S. The Importance of Being Tyrosine: Lessons in Molecular Recognition from Minimalist Synthetic Binding Proteins, Acs Chemical Biology 4, 325–334 (2009).

25. Miersch, S., and Sidhu, S. S. Synthetic antibodies: Concepts, potential and practical considerations, Methods 57, 486–498 (2012).

26. Sidhu, S. S., and Kossiakoff, A. A. Exploring and designing protein function with restricted diversity, Current Opinion in Chemical Biology 11, 347–354 (2007).

27. Akbar, R., Robert, P. A., Pavlović, M., Jeliazkov, J. R., Snapkov, I., Slabodkin, A., Weber, C. R., Scheffer, L., Miho, E., Haff, I. H., et al. A compact vocabulary of paratope-epitope interactions enables predictability of antibody-antigen binding, Cell Reports 34, 108856 (2021).

28. Ferdous, S., and Martin, A. C. R. AbDb: antibody structure database-a database of PDB-derived antibody structures, Database-the Journal of Biological Databases and Curation (2018).

29. Morrison, K. L., and Weiss, G. A. Combinatorial alanine-scanning, Curr Opin Chem Biol 5, 302–307 (2001).

30. Ikeda, J. I., Oda, T., Inoue, M., Uekita, T., Sakai, R., Okumura, M., Aozasa, K., and Morii, E. Expression of CUB domain containing protein (CDCP1) is correlated with prognosis and survival of patients with adenocarcinoma of lung, Cancer Science 100, 429–433 (2009).

31. Scherl-Mostageer, M., Sommergruber, W., Abseher, R., Hauptmann, R., Ambros, P., and Schweifer, N. Identification of a novel gene, CDCP1, overexpressed in human colorectal cancer, Oncogene 20, 4402–4408 (2001).

32. Khan, T., Kryza, T., Lyons, N. J., He, Y., and Hooper, J. D. The CDCP1 Signaling Hub: A Target for Cancer Detection and Therapeutic Intervention, Cancer Research 81, 2259–2269 (2021).

33. Martinko, A. J., Truillet, C., Julien, O., Diaz, J. E., Horlbeck, M. A., Whiteley, G., Blonder, J., Weissman, J. S., Bandyopadhyay, S., Evans, M. J., et al. Targeting RAS-driven human cancer cells with antibodies to upregulated and essential cell-surface proteins, eLife 7, e31098 (2018).

34. Hornsby, M., Paduch, M., Miersch, S., Sääf, A., Matsuguchi, T., Lee, B., Wypisniak, K., Doak, A., King, D., Usatyuk, S., et al. A High Through-put Platform for Recombinant Antibodies to Folded Proteins*, Molecular & Cellular Proteomics 14, 2833–2847 (2015).

35. Hill, Z. B., Martinko, A. J., Nguyen, D. P., and Wells, J. A. Human antibody-based chemically induced dimerizers for cell therapeutic applications, Nature Chemical Biology 14, 112–117 (2018).

36. Casar, B., He, Y., Iconomou, M., Hooper, J. D., Quigley, J. P., and Deryugina, E. I. Blocking of CDCP1 cleavage in vivo prevents Akt-dependent survival and inhibits metastatic colonization through PARP1-mediated apoptosis of cancer cells, Oncogene 31, 3924–3938 (2012).

37. Vandooren, J., Van den Steen, P. E., and Opdenakker, G. Biochemistry and molecular biology of gelatinase B or matrix metalloproteinase-9 (MMP-9): The next decade, Critical Reviews in Biochemistry and Molecular Biology 48, 222–272 (2013).

38. Ra, H.-J., and Parks, W. C. Control of matrix metalloproteinase catalytic activity, Matrix Biology 26, 587–596 (2007).

39. Appleby, T. C., Greenstein, A. E., Hung, M., Liclican, A., Velasquez, M., Villasenor, A. G., Wang, R., Wong, M. H., Liu, X. H., Papalia, G. A., et al. Biochemical characterization and structure determination of a potent, selective antibody inhibitor of human MMP9, Journal of Biological Chemistry 292, 6810–6820 (2017).

40. Koshikawa, N., Hoshino, D., Taniguchi, H., Minegishi, T., Tomari, T., Nam, S. O., Aoki, M., Sueta, T., Nakagawa, T., Miyamoto, S., et al. Proteolysis of EphA2 Converts It from a Tumor Suppressor to an Oncoprotein, Cancer Research 75, 3327–3339 (2015).

41. Schaefer, K., Lui, I., Byrnes, J. R., Kang, E., Zhou, J., Weeks, A. M., and Wells, J. A. Direct Identification of Proteolytic Cleavages on Living Cells Using a Glycan-Tethered Peptide Ligase, Acs Central Science 8, 1447–1456 (2022).

42. Sugiyama, N., Gucciardo, E., Tatti, O., Varjosalo, M., Hyytiainen, M., Gstaiger, M., and Lehti, K. EphA2 cleavage by MT1-MMP triggers single cancer cell invasion via homotypic cell repulsion, Journal of Cell Biology 201, 467–484 (2013).

43. Kabsch, W., and Sander, C. Dictionary of protein secondary structure - pattern-recognition of hydrogen-bonded and geometrical features, Biopolymers 22, 2577–2637 (1983).

44. Hornsby, M., Paduch, M., Miersch, S., Saaf, A., Matsuguchi, T., Lee, B., Wypisniak, K., Doak, A., King, D., Usatyuk, S., et al. A High Through-put Platform for Recombinant Antibodies to Folded Proteins, Molecular & Cellular Proteomics 14, 2833–2847 (2015).

45. Simossis, V. A., and Heringa, J. PRALINE: a multiple sequence alignment toolbox that integrates homology-extended and secondary structure information, Nucleic Acids Res 33, W289–294 (2005).

